# Genome downsizing, physiological novelty, and the global dominance of flowering plants

**DOI:** 10.1101/174615

**Authors:** Kevin A. Simonin, Adam B. Roddy

## Abstract

During the Cretaceous (145-66 Ma), early angiosperms rapidly diversified, eventually outcompeting the ferns and gymnosperms previously dominating most ecosystems. Heightened competitive abilities of angiosperms are often attributed to higher rates of transpiration facilitating faster growth. This hypothesis does not explain how angiosperms were able to develop leaves with smaller, but densely packed stomata and highly branched venation networks needed to support increased gas exchange rates. Although genome duplication and reorganization have likely facilitated angiosperm diversification, here we show that genome downsizing facilitated reductions in cell size necessary to construct leaves with a high density stomata and veins. Rapid genome downsizing during the early Cretaceous allowed angiosperms to push the frontiers of anatomical trait space. In contrast, during the same time period ferns and gymnosperms exhibited no such changes in genome size, stomatal size, or vein density. Further reinforcing the effect of genome downsizing on increased gas exchange rates, we found that species employing water-loss limiting crassulacean acid metabolism (CAM) photosynthesis, have significantly larger genomes than C3 and C4 species. By directly affecting cell size and gas exchange capacity, genome downsizing brought actual primary productivity closer to its maximum potential. These results suggest species with small genomes, exhibiting a larger range of final cell size, can more finely tune their leaf physiology to environmental conditions and inhabit a broader range of habitats.

## Introduction

The abrupt origin and rapid diversification of the flowering plants during the mid-Cretaceous, and their eventual dominance globally, has long been considered an ‘abominable mystery’ ^1^. While the cause of their high diversity has been attributed primarily to coevolution with pollinators and herbivores, many hypotheses have been posed to explain why angiosperms were able to become ecologically dominant in most terrestrial ecosystems. A common theme among these hypotheses has been the idea that angiosperms developed a set of physiological traits that allowed them to achieve higher rates of primary productivity than either the ferns or the gymnosperms ^2^. Terrestrial primary productivity is determined by the photosynthetic capacity of leaves, and one of the greatest biophysical limitations to photosynthetic rates across all the major clades of terrestrial plants is the leaf surface conductance to CO_2_ and water vapor. In order for CO_2_ to diffuse from the atmosphere into the leaf, the wet internal surfaces of leaves must be exposed to the dry ambient atmosphere, which can cause leaf desiccation and prevent further CO_2_ uptake. As a consequence, increasing leaf surface conductance to CO_2_ also requires increasing rates of leaf water transport in order to avoid desiccation^3^.

Both theory and empirical data suggest that among all major clades of terrestrial plants the upper limit of leaf surface conductance to CO_2_ and water vapor is tightly coupled to biophysical limitations on cell size ^4-7^. Cellular allometry, in particular the scaling of genome size, nuclear volume, and cell size represents a direct physical constraint on the number of cells that can occupy a given space and, as a result, on the distance between cell types and tissues ^8^. Because leaves with many small stomata and a high density of veins promote higher rates of gas exchange than leaves with fewer, larger stomata and larger, less dense veins ^9^, variation in cell size can drive large changes in potential carbon gain ^10^. Without reducing cell size, increasing stomatal and vein densities would displace other important tissues, such as photosynthetic mesophyll cells ^11^. Therefore, the densities of stomata on the leaf surface and of veins inside the leaf are inversely related to the sizes of guard cells and xylem elements of which they are comprised.

While numerous environmental and physiological factors can influence the final sizes of somatic eukaryotic cells, the minimum size of meristematic cells and the rate of their production are strongly constrained by nuclear volume, more commonly measured as genome size ^12-15^. Among land plants, the bulk DNA content of cells varies by three orders of magnitude, with the angiosperms exhibiting both the largest range in genome size and the smallest absolute genome sizes ^16^. Whole-genome duplications and subsequent genomic rearrangements rapidly change genome size and are thought to have directly contributed to the unparalleled diversity in anatomical, morphological, and physiological traits of the angiosperms ^15,17-21^. We extend this prior work and predict that genome size variation is not only responsible for gene diversification but also directly controls minimum cell size and, thus, is the underlying variable directly influencing both stomatal size and density and leaf vein density thus directly influencing rates of leaf gas exchange across the major clades of terrestrial plants.

To test whether genome downsizing among the angiosperms drove the anatomical and physiological innovations that resulted in their ecological dominance over other major clades of terrestrial plants, we compiled data for genome size, cell size (guard cell length, *l*_*g*_), leaf vein density (*D*_*v*_), and maximum and operational leaf surface conductance to CO_2_ and water vapor (*g*_*s,max*_ and *g*_*s,op*_, respectively) for almost 1100 species of ferns, gymnosperms, and angiosperms. If genome downsizing were critical for angiosperm success, then we expect genome size to have declined rapidly during early angiosperm evolution but, perhaps, to have remained unchanged among the ferns and gymnosperms. Furthermore, if genome size constrains *l*_g_ and *D*_v_, then evolutionary changes in genome size should precede changes in both *l*_g_ and *D*_v_. Finally, we predict that the benefits of genome downsizing on carbon gain should be greatest when photosynthesis and transpiration are proportional, such as in plant species that possess C3 and C4 photosynthetic metabolism, in contrast to species employing crassulacean acid metabolism (CAM), which decouples gas exchange from periods of high evaporative demand. If these predictions about the biophysical effects of genome size and its evolution are supported, then genome downsizing among the angiosperms led directly to their greater potential and realized primary productivity, contributing to their rapid domination of ecosystems globally.

## Results

### Trait correlations (genome vs vein density and cell size) and phylogenetic independent contrasts

Genome size varied substantially among major clades (Figure 1) and was a strong predictor of anatomical traits across the major groups of terrestrial plants even when accounting for phylogeny. Genome size explained 42% of between species variation in *l*_g_ across the major groups of terrestrial plants (Figure 2a). Additionally, a single relationship predicted *l*_g_ from genome size across all species. Similarly, a strong negative correlation existed between genome size and both *D*_s_ (Figure 2b; *R*^*2*^ = 0.32) and *D*v (Figure 2c; *R*^*2*^ = 0.46). Among major clades and within the angiosperms, traits showed strong, significant correlations between PICs, highlighting the coordinated evolution of these traits repeatedly throughout the history of seed plants (Table S2).

**Figure 1.**
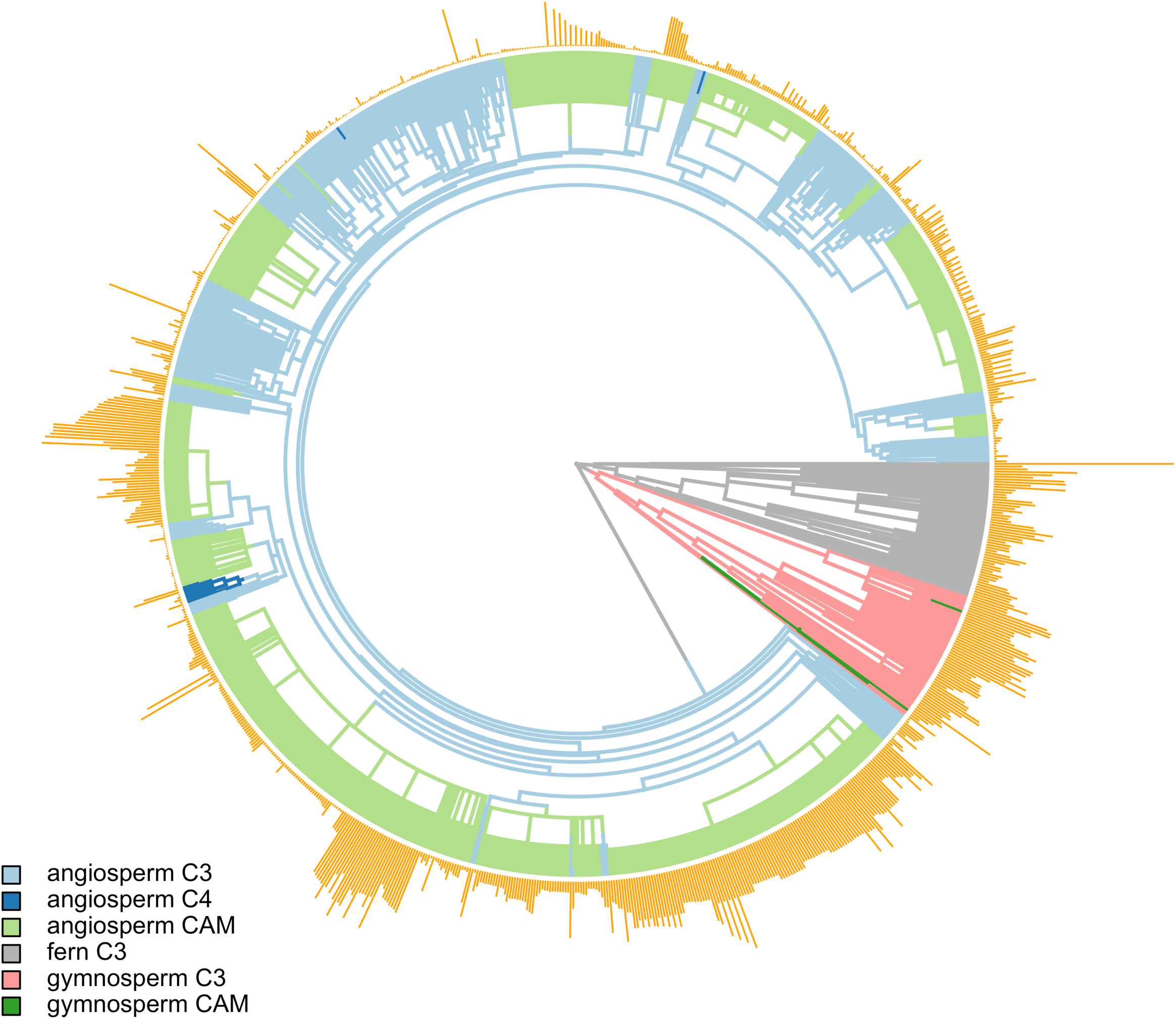
The distribution of genome size among 1035 land plants. The family level phylogeny has branches colored according to one random stochastic character map of photosynthetic pathways (C3, C4, CAM) among clades (ferns, gymnosperms, angiosperms). Orange bars at the tips are scaled proportional to genome size for each terminal species.

**Figure 2.**
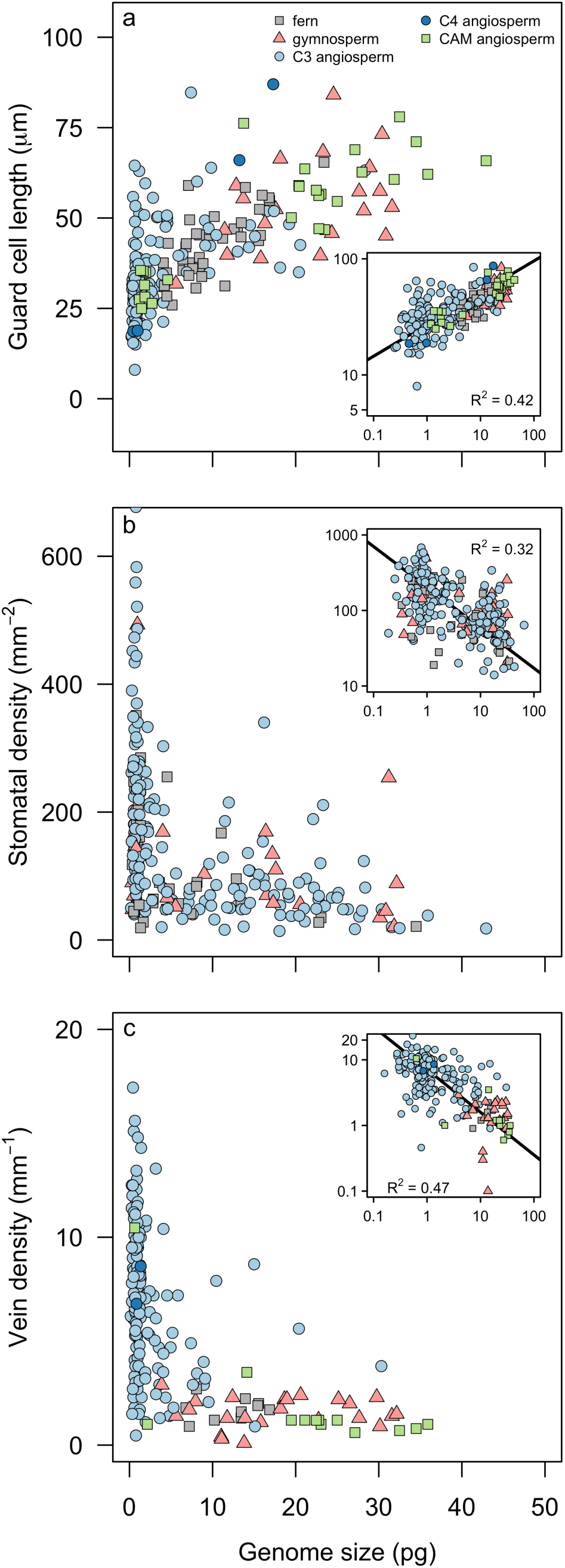
Relationships between genome size and anatomical traits: (a) *l*_*g*_, (b) *D*_*s*_, and (c) *D*_*v*_. In all panels, insets show log-log relationships and *R*^2^ values are from standard major axis regressions. Correlations for phylogenetically corrected relationships are in Table S2.

### Biophysical scaling relationships: maximum and operational leaf surface conductance

Within clades, only the angiosperms exhibited a significant relationship between genome size and either *g*_s, max_ or *g*_s, op_. Sample sizes among the ferns and gymnosperms for these traits were quite low, precluding statistical significance. Yet, ferns and gymnosperms fell within the ranges of *g*_s, max_ and *g*_s, op_ defined by the angiosperms. The global scaling relationships among all species between genome size and either *g*_s,max_ or *g*_s,op_ were not significantly different than those for only the angiosperms, suggesting that a single relationship may exist between genome size and stomatal conductance (Figure 3).

**Figure 3.**
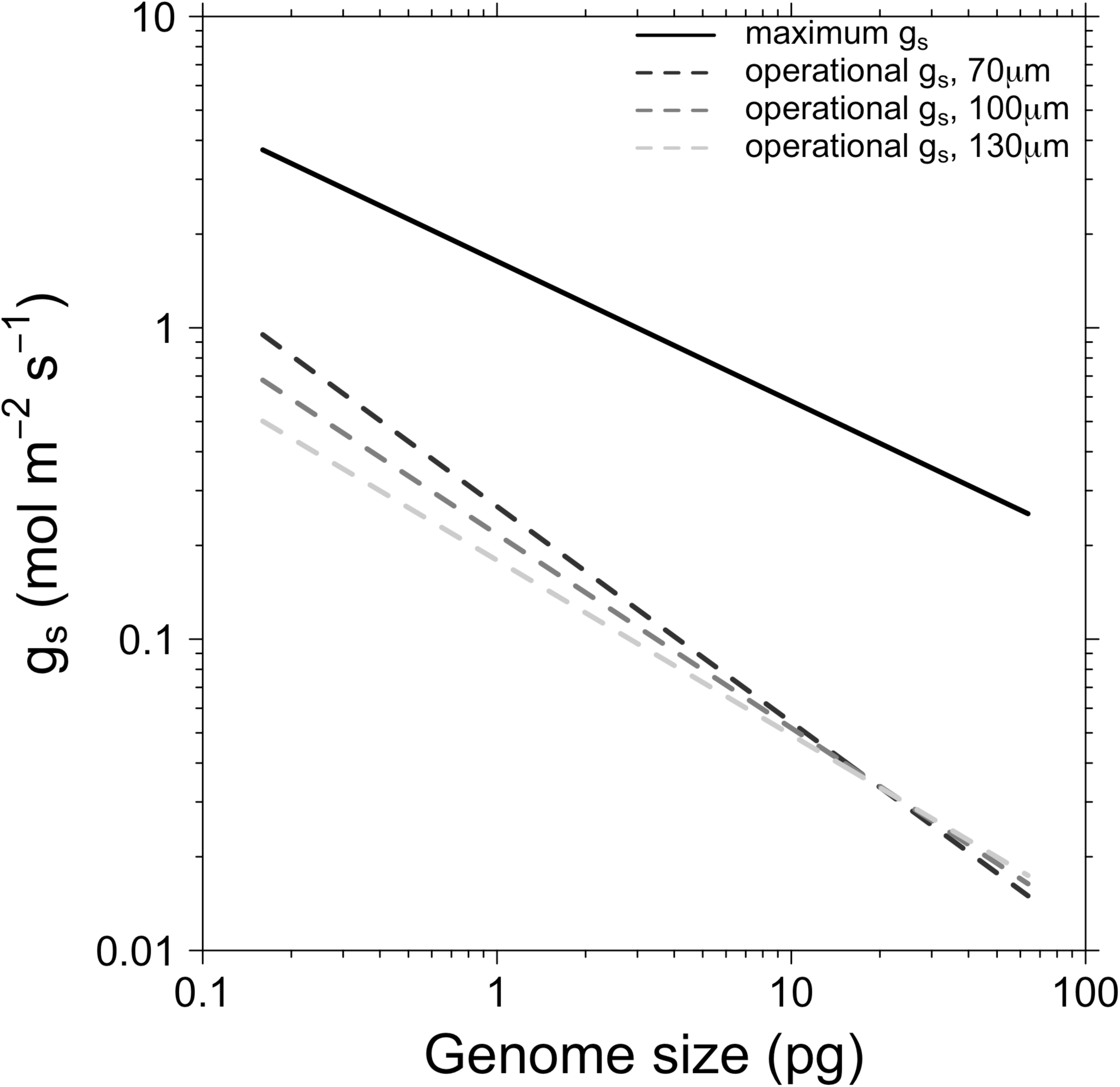
The relationships between genome size and maximum (solid line; *R*^2^ = 0.25) and operational (dashed lines) stomatal conductance, plotted on a log-log scale. Operational stomatal conductance was modeled under assumptions of three leaf thicknesses (70 μm, *R*^2^ = 0.46; 100 μm, *R*^2^ = 0.44; 130 μm, *R*^2^ = 0.43). Points are omitted for clarity. Correlations for phylogenetically corrected relationships are in Table S2.

Regardless of the leaf thickness (70, 100, 130 *μ*m) used to calculate *g*_s,op_, the scaling relationships between genome size and *g*_s,op_ were significantly steeper than the relationship between genome size and *g*_s,max_ (all P < 0.001). Therefore, across species, shrinking the genome brings *g*_s,op_ closer to *g*_s,max_ (Figure 3, Table 1).

**Table 1.** Standard major axis regressions of *D*_*v*_, *l*_*g*_, *g*_s,max_, and *g*_s,op_ versus genome size for all species and for each clade separately. Asterisks indicate significance level: *P < 0.05; **P < 0.01; ***P < 0.001

|  | all |  |  | angiosperm |  |  | gymnosperm |  |  | fern |  |  |
| --- | --- | --- | --- | --- | --- | --- | --- | --- | --- | --- | --- | --- |
|  | slope | intercept | R<sup>2</sup> | slope | intercept | R<sup>2</sup> | slope | intercept | R<sup>2</sup> | slope | intercept | R<sup>2</sup> |
| $D_v$ | -0.646<br>(-0.716,<br>-0.583) | 0.848<br>(0.799,<br>0.897) | 0.463<br>*** | -0.641<br>(-0.727,<br>-0.566) | 0.838<br>(0.789,<br>0.887) | 0.329<br>*** | 1.34 (0.864,<br>2.078) | -1.488<br>(-2.244,<br>-0.732) | 0.002 | 0.940<br>(0.454,<br>1.944) | -0.778<br>(-1.580,<br>0.025) | 0.054 |
| $l_g$ | 0.274<br>(0.249,<br>0.301) | 1.435<br>(1.414,<br>1.456) | 0.420<br>*** | 0.300<br>(0.270, 0.333) | 1.433<br>(1.411,<br>1.454) | 0.477<br>*** | 0.529<br>(0.354,0.792) | 1.036<br>(0.749,<br>1.324) | 0.300<br>* | 0.480<br>(0.383,<br>0.602) | 1.167<br>(1.060,<br>1.274) | 0.523<br>*** |
| $g_{s,max}$ | -0.449<br>(-0.509,<br>-0.397) | 0.214<br>(0.167,<br>0.260) | 0.245<br>*** | -0.447<br>(-0.514,<br>-0.388) | 0.208<br>(0.160,<br>0.255) | 0.170<br>*** | 1.178 (0.668,<br>2.078) | -1.750<br>(-2.646, -<br>0.854) | 0.004 | 0.971<br>(0.425,<br>2.221) | -1.201<br>(-2.11,<br>-0.288) | 0.352 |
| $g_{s,op}$<br>70µm | -0.693<br>(-0.769,<br>-0.625) | -0.573<br>(-0.626,<br>-0.522) | 0.457<br>*** | -0.662<br>(-0.750,<br>-0.584) | -0.586 (-<br>0.635, -<br>0.536) | 0.335<br>*** | 1.605 (1.010,<br>2.551) | -3.295<br>(-4.252, -<br>2.338) | 0.001 | 1.169<br>(0.566,<br>2.415) | -2.473<br>(-3.468,<br>-1.479) | 0.060 |

### Trait evolution through time

Compared to the ferns and gymnosperms, genome sizes, *D*_v_, and *l*_g_ of the angiosperms all evolved into new regions of trait space during the Cretaceous (Figure 4), increasing rates of carbon assimilation and ushering in more rapidly growing forests. For all three traits, the logarithmic curve fit the extreme values better than a linear relationship (genome size **Δ**AIC = 12.89; *D*_v_ **Δ**AIC = 11.43; *l*_g_ **Δ**AIC = 24.69). In contrast to the angiosperms, fern and gymnosperm lineages exhibited no such change in any of the traits during the Cretaceous. For fern and gymnosperm traits, the linear fit including a slope and intercept was not significantly better than the model lacking a slope (i.e. the mean reconstructed trait value), except for fern minimum *l*_g_, which was better modeled by a linear regression with a slope, although this model indicated that fern *l*_g_ increased through time (Figure 4).

**Figure 4.**
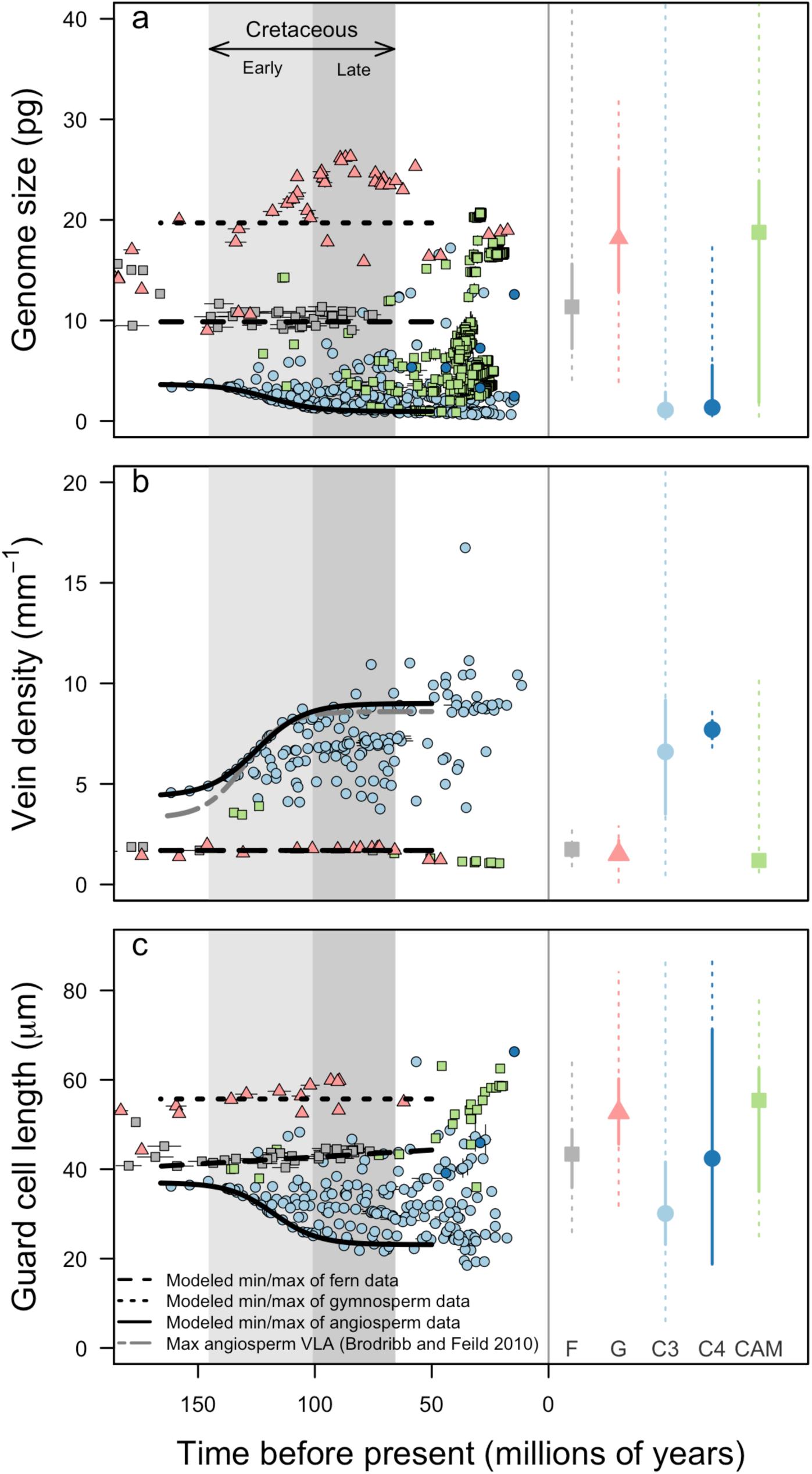
Ancestral state reconstructions of genome size, vein density (*D*_*v*_), and guard cell length (*l*_*g*_) through time for angiosperms (colored circles), gymnosperms (grey triangles), and ferns (grey squares). Error bars around reconstructed values represent error due to phylogenetic uncertainty. The shaded timespan indicates the Cretaceous, during which most major lineages of angiosperms diversified. Lines represent the best-fit models through the lower (genome size and *l*_*g*_) and upper (*D*_*v*_) 10% of reconstructed values. (a) Genome size was unchanged during the Cretaceous for the ferns (genome size = 9.91, df = 2, P < 0.001) and the gymnosperms (genome size = 19.62, df = 2, P < 0.001), while minimum genome size among the angiosperms decreased rapidly during the Cretaceous (genome size = 0.99 + 3.06/(1 + e^(-time – 120.74)/9.03; df = 5, P < 0.001). (b) Similar to the results of Brodribb and Field (2010), the upper limit of reconstructed *D*_*v*_ through time increased significantly for the angiosperms (*D*_*v*_ = 3.93 + 5.29/(1 + e^(-(time-124.47)/(-13.49)); df = 5, P < 0.001). However, vein densities of fern (*D*_*v*_ = 1.70; df = 2, P < 0.01) and gymnosperm lineages (*D*_*v*_ = 1.68; df = 2, P < 0.001) remained unchanged during the same time period. (c) Similarly, *l*_*g*_ declined rapidly among angiosperms (*l*_*g*_ = 23.64 + 13.82/(1 + e^(-(time −118.48)/9.31); df = 5, P < 0.001), while *l*_*g*_ of ferns (*l*_*g*_ = 42.52, df = 2, P < 0.001) and gymnosperms (*l*_*g*_ = 55.66, df = 2, P < 0.001) remained unchanged during the Cretaceous. Marginal plots on the right represent the median (points), interquartile ranges (solid lines) and ranges (dotted lines) of extant trait values. Angiosperm data have been plotted separately for species exhibiting each photosynthetic pathway. The two CAM gymnosperms were included with C3 gymnosperms in these analyses.

Genome size evolution among C3 species was best modeled by allowing for different rates of trait evolution for the three clades, consistent with our prediction that the angiosperms capitalized on genome downsizing. Although we had predicted that OU models, which model stabilizing selection around optimum trait values, would best fit the data, the best-fitting model was instead the Brownian motion model that included different rates for each clade. In this model, the rate parameter indicates the standard deviation of trait values around the phylogenetic mean; thus a faster rate is indicative of greater trait variance. In all 100 simulations, the Brownian motion model provided the best fit with **Δ**AIC = 14.33 ± 0.17, compared to the second-best fitting model in each iteration (Table 2). Across all C3 taxa, genome size evolved faster in the ferns (0.19 ± 0.0009) and gymnosperms (0.14 ± 0.0008) than in the angiosperms (0.088 ± 0.0006). Similarly, in the combined analysis that incorporated all clades and photosynthetic pathways, a Brownian motion model with multiple rates best described the data in all 100 simulations (**Δ**AIC = 130.14 ± 1.20), and the modeled parameters were similar to those from the other models (Table 2).

**Table 2.**
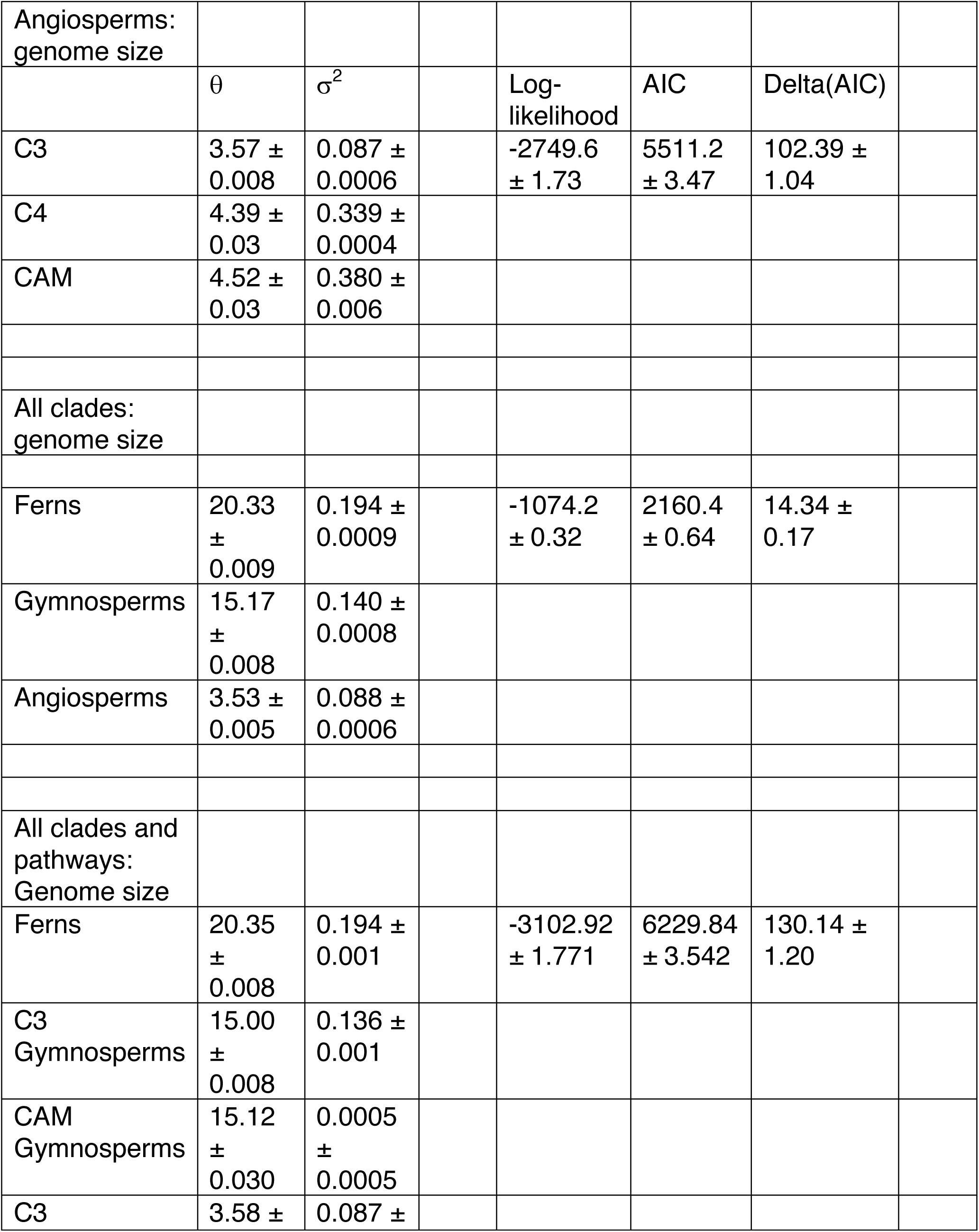
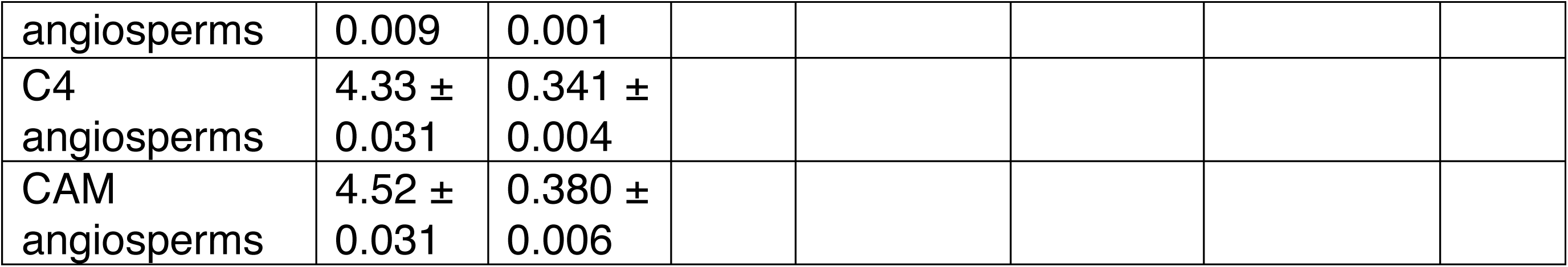
Univariate evolutionary modeling of genome size was best fit by a Brownian motion model with multiple rates. Parameter values are means ± standard error of 100 replicate simulations accounting for phylogenetic uncertainty. θ = genome size at the phylogenetic root, σ^2^= rate of evolution.

|  |  |  |  |  |  |  |  |
| --- | --- | --- | --- | --- | --- | --- | --- |
| Angiosperms:<br>genome size |  |  |  |  |  |  |  |
| | $\theta$ | $\sigma^2$ | | Log-likelihood | AIC | Delta(AIC) | |
| C3 | 3.57 ± 0.008 | 0.087 ± 0.0006 | | -2749.6 ± 1.73 | 5511.2 ± 3.47 | 102.39 ± 1.04 | |
| C4 | 4.39 ± 0.03 | 0.339 ± 0.0004 | | | | | |
| CAM | 4.52 ± 0.03 | 0.380 ± 0.006 | | | | | |
| All clades:<br>genome size |  |  |  |  |  |  |  |
| Ferns | 20.33 ± 0.009 | 0.194 ± 0.0009 | | -1074.2 ± 0.32 | 2160.4 ± 0.64 | 14.34 ± 0.17 | |
| Gymnosperms | 15.17 ± 0.008 | 0.140 ± 0.0008 | | | | | |
| Angiosperms | 3.53 ± 0.005 | 0.088 ± 0.0006 | | | | | |
| All clades and<br>pathways:<br>Genome size |  |  |  |  |  |  |  |
| Ferns | 20.35 ± 0.008 | 0.194 ± 0.001 | | -3102.92 ± 1.771 | 6229.84 ± 3.542 | 130.14 ± 1.20 | |
| C3<br>Gymnosperms | 15.00 ± 0.008 | 0.136 ± 0.001 | | | | | |
| CAM<br>Gymnosperms | 15.12 ± 0.030 | 0.0005 ± 0.0005 | | | | | |
| C3 | 3.58 ± | 0.087 ± | | | | | |

|  |  |  |
| --- | --- | --- |
| angiosperms | 0.009 | 0.001 |
| C4<br>angiosperms | 4.33 ±<br>0.031 | 0.341 ±<br>0.004 |
| CAM<br>angiosperms | 4.52 ±<br>0.031 | 0.380 ±<br>0.006 |

Among the angiosperms, genome size evolution differed with photosynthetic pathway, reflecting that genome size-cell size allometry imposes different constraints on C3 and CAM species (Figures 1, 5). First, genome size evolution was best modeled by a Brownian motion process (100 out of 100 simulations, **Δ**AIC = 102.39 ±1.04; Table 2) that allowed for multiple rates of evolution for lineages employing the different photosynthetic pathways. CAM lineages had the largest estimated phylogenetic mean genome size (equivalent to the estimated ancestral genome size) and also the fastest rate of genome size evolution, in both balanced (phylogenetic mean: *t* = 18.61, df = 105.11, P < 0.0001; rate: *t* = 35.36, df = 99.65, P < 0.0001) and unbalanced (phylogenetic mean: *t* = 30.89, df = 113.66, P < 0.0001; rate: *t* = 47.53, df = 100.99, P < 0.0001) species sampling (Figure 4). Second, we tested whether there were time lags between shifts in genome size and shifts in either *D*_v_ or *l*_g_ associated with transitions between photosynthetic pathways. If genome size fundamentally constrains *D*_v_ and *l*_g_, then shifts in genome size should either coincide with or precede shifts in the other traits, but genome size should not lag behind either *D*_v_ or *l*_g_. Although in 96 of 100 simulations vein density lagged behind genome size, support for this model was weak (ΔAIC = 2.80 ± 0.16), suggesting that there has been little or no lag between *D*_v_ and genome size. Similarly, although shifts in *l*_g_ lagged behind shifts in genome size in 80 of 100 simulations, support was weak (ΔAIC = 0.99 ± 0.09). In the other 20 simulations, there was no lag between shifts in genome size and *l*_g_ (ΔAIC = 1.32 ± 0.45), further suggesting that genome size and cell size evolve in unison. In none of the time lag simulations did genome size lag behind either *D*_v_ or *l*_g_, strengthening support for genome size fundamentally constraining both *l*_g_ and *D*_v_.

**Figure 5.**
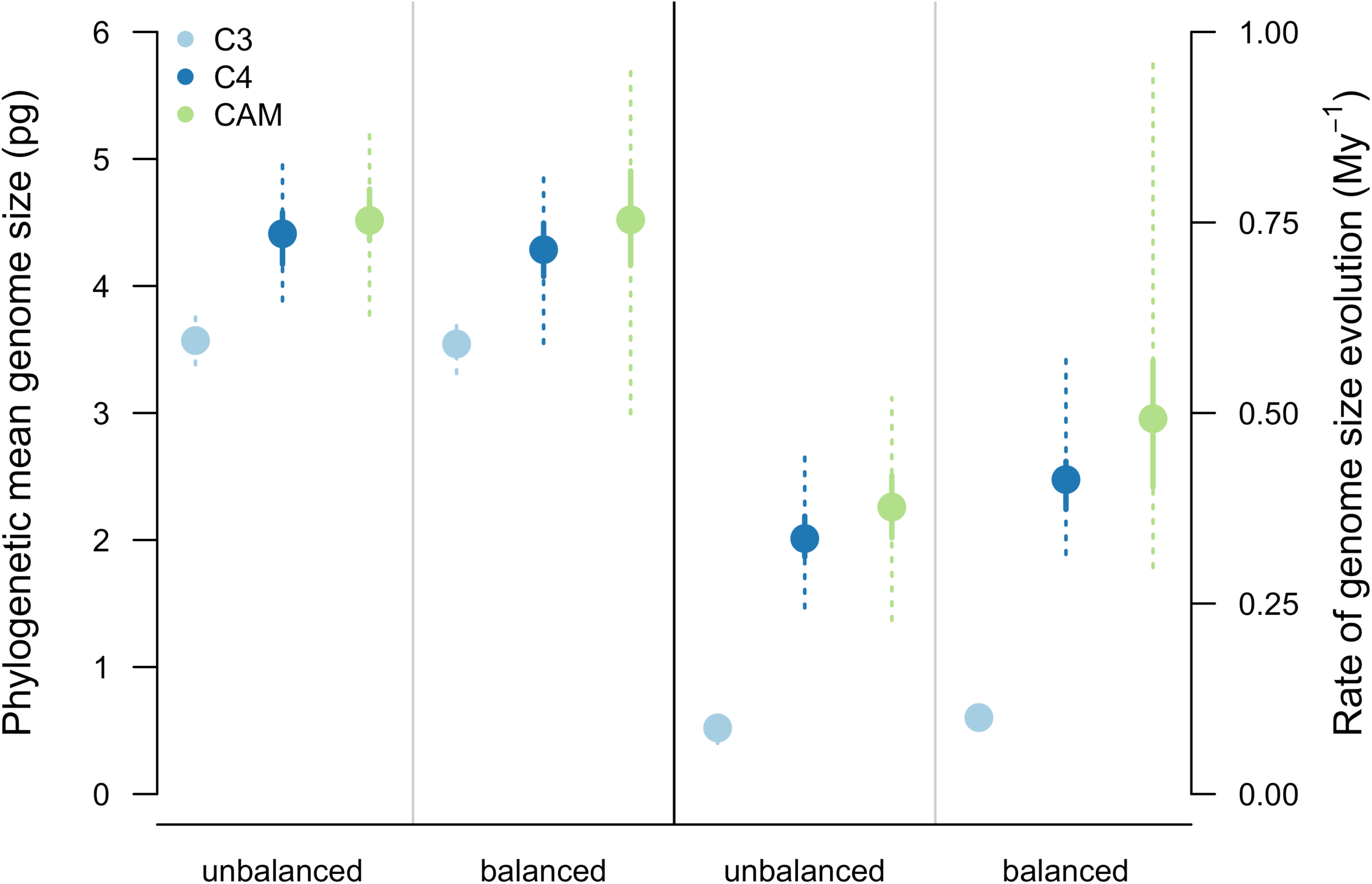
Differences in reconstructed genome size and the rate of genome size evolution for angiosperms differing in photosynthetic pathway. The reconstructed genome size and the rate of genome size evolution differed among C3 and CAM species, regardless of whether equivalent numbers of C3 and CAM species (‘balanced’, 271 species each) were randomly sampled or whether all CAM species were included in the analysis (‘unbalanced’). The rate of genome size evolution is the Brownian motion rate parameter calculated separately for each photosynthetic pathway. Points are medians, solid lines are interquartile ranges, and dotted lines are ranges of modeled parameters.

## Discussion

Our results suggest that the basis for developing leaves with the potential for high rates of gas exchange derive not exclusively from common developmental programs nor from genetic correlations (i.e. linkage between genes controlling both traits), but, even more fundamentally, from biophysical scaling constraints that limit minimum cell size ^4,38^. These scaling relationships between genome size and gas exchange rates as well as analyses of trait evolution suggest that genome downsizing among the angiosperms permitted the evolution of the anatomical traits responsible for increased rates of photosynthesis and biomass accumulation (Figures 2-4). Importantly, while genome downsizing has been critical to increasing leaf gas exchange rates among the angiosperms, it was not a key innovation that occurred only at the root of the angiosperm phylogeny. Rather, the angiosperms exhibit a wide range of genome sizes, and coordinated changes in genome size and physiological traits have repeatedly occurred throughout the evolutionary history of the angiosperms (Table S2). Whole-genome duplications have been particularly important in promoting diversification among the angiosperms ^17^ yet result in larger, physiologically detrimental, genomes. Our results suggest that genome downsizing is critical to recovering leaf gas exchange capacity subsequent to genome duplications.

The ecological revolution ushered in by the angiosperms is due largely to the biophysical benefits associated with decreasing genome and cell sizes. If heightened competitive ability among the angiosperms drove their ecological dominance, then innovations that allowed minimum cell size to decline were critical to this transformative process ^38^. Because genome size provides a boundary on minimum cell size, genome size has numerous consequences for the structure and organization of cells and tissues in leaves, which directly influence metabolic rates. Specifically, unlike ferns and gymnosperms, angiosperms were able to develop leaves with numerous, small stomata and a high density of veins because of rapid reductions in genome size during the Cretaceous (Figures 2, 4). Non-angiosperm lineages exhibited no similar changes in these traits during the same time, despite a single, universal scaling relationship in all major clades of terrestrial plants between genome size and anatomical (*D*_v_ and *l*_g_) and physiological (*g*_s,max_ and *g*_s,op_) traits. Across seed plants, genome downsizing effectively brings actual productivity closer to its theoretical maximum (Figure 3), allowing the angiosperms to outcompete other land plants.

Cell size has direct and predictable effects on gas diffusion across the leaf epidermis, and, as we show here, also on the supply of liquid water to the leaf. Physical resistance to diffusion across leaf surfaces is ultimately determined by the size of epidermal cells, and the maximum diffusive conductance of CO_2_ and water vapor (*g*_s, max_) is higher in leaves with numerous, small stomata ^4,6,7^. While the effects of cell size on leaf epidermal properties have been well characterized, the effects of cell size on the efficiency of liquid water supply through the leaf are, perhaps, less obvious. Given a constant leaf volume, increasing *D*_v_ without displacing photosynthetic mesophyll cells requires reductions in vein and conduit sizes that can only be accomplished by decreasing cell size ^11,39^. However, smaller conduits have higher hydraulic resistances. To overcome the increase in resistance associated with reducing conduit sizes, other innovations in xylem anatomy that reduce hydraulic resistance have been hypothesized to facilitate narrower xylem conduits and high *D*_v_. In particular, the development of low resistance end walls between adjacent cells is thought to have given angiosperms a hydraulic advantage as conduit diameters decreased. Only in angiosperm lineages with very high *D*_v_ do primary xylem have simple perforation plates, which have lower resistance to water flow than scalariform perforation plates ^11^. Similarly, the low resistance of gymnosperm torus-margo pits compared to angiosperm pits can result in higher xylem specific hydraulic conductivity for small diameter conduits ^40^. In both cases, while smaller conduits have higher resistance, this potential cost has been offset by other innovations that reduce hydraulic resistance at the scale of the whole xylem network. The requirement of these other changes to xylem anatomy to occur before potential gains from reduced conduit sizes can be realized may explain why evolutionary shifts in *D*_v_ almost always lagged behind shifts in genome size associated with transitions between photosynthetic pathways. In contrast, shifts in *l*_g_ were less likely to lag behind shifts in genome size, instead evolving concurrently with genome size, probably due to the direct and simple effect of genome size on *l*_g_ without the need for other traits to evolve.

While genome size limits minimum cell size, final cell size can vary greatly as cells grow and differentiate. After cell division and during cell expansion, various factors influence how large a cell becomes. Intracellular turgor pressure overcomes the mechanical rigidity of the cell wall to enlarge cellular boundaries. The magnitude of turgor pressure is itself controlled by water availability around the cell and the osmotic potential inside the cell. Final cell size is controlled therefore by both biotic and abiotic factors that influence pressure gradients in and around the cell. By reducing the lower limit of cell size, genome downsizing expands the range of final cell size that is possible. Thus, species that can vary cell size across a wider range can more finely tune their leaf anatomy to match environmental constraints on leaf gas exchange. Indeed, *D*_v_, *l*_g_, and stomatal conductance are more variable among species with small genomes, and the variance in these traits unexplained by genome size is likely due to environmental variation (Figures 2-4), although analyses of intraspecific genome size variation are needed to further clarify the potential links between genome size variation and environmental variation. Interestingly, only the angiosperms occupy this region of trait space, and the angiosperms tend to be more productive than either the ferns or the gymnosperms across a broad range of environmental conditions. Furthermore, genome size may predict ecological breadth even within species insofar as species with small genomes can exhibit greater plasticity in final cell size and inhabit a wider range of environmental conditions. Thus, rapid genome downsizing by the angiosperms during the Cretaceous likely explains not only their greater potential and realized primary productivity (Figure 3) but also why they were able to expand into and create new ecological habitats, fundamentally altering the global biosphere and atmosphere ^41^.

Yet, not all angiosperms have small genomes (Figures 1-2). Genome size-cell size allometry determines physiological function within a given environment when photosynthetic rates are proportional to transpiration rates. This is certainly the case for species employing the C3 and C4 photosynthetic pathways. However, due to higher water use efficiency of C4 photosynthesis, the physiological effects of genome size variation may be slightly weaker in C4 species than they are in C3 species, as reflected in the slightly larger genomes and guard cell lengths of C4 species. Nonetheless, *D*_v_ and photosynthetic metabolism in many C4 plants are intimately linked due to the physical arrangement of the sites of carboxylation into a layer of cells surrounding the veins (i.e. bundle sheath cells). Because of this physical association, increasing the number of cells that are able to assimilate CO_2_ requires an increase in the number of veins. In contrast to both C3 and C4 photosynthetic metabolism, species employing CAM photosynthesis effectively decouple carbon uptake from periods of relatively high evaporative demand. For CAM species, the constraints of genome size on the coordination between carbon gain and water loss are minimal, and, as a result, CAM species have significantly larger genomes than either C3 or C4 species (Figures 4-5). CAM lineages also have faster rates of genome size evolution than C3 lineages, suggesting that genome size may be more constrained in C3 lineages because cell size has a direct and substantial effect on gas exchange rates (Figure 5). However, the limited taxonomic resolution of CAM photosynthesis may be biasing our estimates of evolutionary rates. There are undoubtedly C3 species and C3-CAM intermediates that we have classified as strictly CAM, which would increase the disparity in genome size and lead to an underestimation of the difference in genome size between C3 and CAM species but an overestimation of the rate of genome size evolution among species classified as strictly CAM. Nonetheless, the difference in genome size between extant C3 and CAM species highlights the fundamental importance of genome downsizing in raising the limits of leaf gas exchange when carbon uptake is directly coupled to water loss. By no means should this trivialize the ecological importance of CAM photosynthesis. Rather, it reinforces the innovativeness of the angiosperms because genome downsizing is not the only strategy conferring ecological success; CAM photosynthesis, regardless of genome size, has allowed colonization of marginal environments often uninhabitable by C3 species.

## Conclusion

The rapid diversification and spread of angiosperms during the Cretaceous dramatically restructured terrestrial ecosystems ^41,42^. While their heightened diversification rates have long been thought to result from a combination of unique traits that allowed them to coevolve with pollinators and herbivores ^43-47^, only recently have hypotheses about how angiosperms became ecologically dominant been considered. Central to these hypotheses has been that the angiosperms became competitively more successful due to faster growth rates ^48^, supported by higher rates of photosynthesis and transpiration ^9,30,42^. Anatomical innovations that appeared among the angiosperms–smaller, more abundant stomata and narrower, more densely packed leaf veins–that support higher rates of transpiration and photosynthesis would have been particularly advantageous as atmospheric CO_2_ concentration declined during the Cretaceous. These traits are unique to the angiosperms and due, we show, to reductions in cell and genome sizes that occurred after the appearance of early angiosperms. Smaller genomes and cells increased leaf surface conductance to CO_2_ and enabled higher potential and realized primary productivity. Interestingly, the physiological benefits of small genomes and cells are realized only when photosynthetic rates are proportional to transpiration rates; species employing CAM photosynthesis avoid assimilating CO_2_ during periods of high evaporative demand driven by light interception. As a result, CAM species can have larger genomes without the physiological costs that C3 species might incur. Additionally, CAM species often inhabit marginal habitats characterized by limited water availability and nutrient cycling that are unable to support high rates of primary productivity ^49^. Furthermore, because genome downsizing lowers the limit of minimum cell size, final cell size can vary much more widely, which facilitates a closer coupling of anatomy and physiology with environmental conditions. Therefore, genome downsizing has increased the range of habitable environments and allowed angiosperms to outcompete other land plants in almost every ecosystem.

## Methods

### Leaf traits

Published data for guard cell length (*l*_g_), stomatal density (*D*_s_), and vein density (*D*_v_) were compiled from the literature (Table S1). Genome size data for each species were taken from the Plant DNA C-values database (release 6.0, December 2012), managed by the Royal Botanic Gardens, Kew ^22^. In total, our dataset comprised 1087 species of vascular plants, of which 979 were angiosperms, 54 were gymnosperms, and 54 were ferns. For the 979 angiosperms in the dataset, there were *D*_v_ data for 164 and guard cell size data for 220. Similarly, there were *D*_v_ data for 23 gymnosperms and for 10 ferns, and there were *l*_g_ data for 20 gymnosperms and for 41 ferns. The large discrepancy between the total number of angiosperms in the dataset and the number of angiosperms with leaf trait data is due to inclusion of genome size data for CAM species that lacked leaf trait data.

Because different photosynthetic pathways (C3, C4, CAM) employ different strategies of maintaining water balance, the effects of genome size-cell size allometry may differ among species employing different photosynthetic pathways. We tested this hypothesis by comparing genome size evolution among the angiosperms. We expected the largest difference to exist between C3 and CAM species, and so we focused the analysis on this comparison. The taxonomic distribution of CAM photosynthesis was based on Smith and Winter ^23^, which provides a list of genera exhibiting CAM photosynthesis. Undoubtedly, some of these genera include C3 species (either C3-CAM intermediates or exclusively C3), which would lead to a conservative estimate of the differences in genome size between C3 and CAM species. Of the 973 angiosperms in this analysis, 271 were C3, nine were C4, and 647 were CAM.

### Calculating maximum and operational stomatal conductance

For each species in our database with anatomical traits, we calculated the maximum stomatal conductance and the operational stomatal conductance. Maximum stomatal conductance (*g*_s, max_) is defined by the dimensions of stomatal pores and their abundance, and represents the biophysical upper limit of gas diffusion through the leaf epidermis. Anatomical measurements of guard cells were used to calculate *g* _s,max_ as ^4,5^:

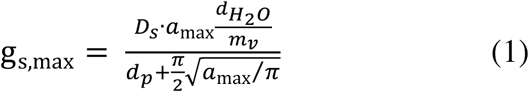

where *d*_H2O_ is the diffusivity of water in air (0.0000249 m^2^ s^-1^), *m*_v_ is the molar volume of air normalized to 25 °C (0.0224 m^3^ mol^-1^), *D*_s_ is stomatal density (mm^-2^), *a*_max_ is maximum stomatal pore size, and *d*_p_ is the depth of the stomatal pore. The *a*_max_ term can be approximated as: π(*l*_p_ /2)^2^, where *l*_p_ is stomatal pore length with *l*_p_ being approximated as *l*_*g*_ /2, where *l*_*g*_ is guard cell length ^4,24^. *d*_p_ is assumed to be equal to guard cell width (*W*). If *W* was not reported *d*_p_ was estimated as 0.36·*l*_g_ ^7^.

Operational stomatal conductance (*g*_s,op_), by contrast, more accurately defines the stomatal conductance leaves attain under natural conditions when limitations in leaf hydraulic supply constrain stomatal conductance. We used an empirical model of g _s, op_ ^3^ that directly relates *D*_v_ to stomatal conductance during periods of steady state transpiration as:

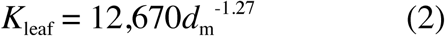

where:

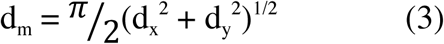

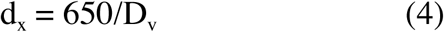

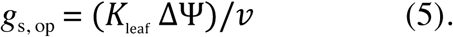

*K*_leaf_ is leaf hydraulic conductance (mmol m^-2^ s^-1^ MPa^-1^), *d*_m_ is the post vein distance to stomata *(μ*m), *d*_x_ is the maximum horizontal distance from vein to the stomata (μm), *d*_y_ is the distance from vein to the epidermis *(μ*m), ΔΨ is the water potential difference between stem and leaf (set to 0.33 MPa ^25^) and *v* is vapor pressure deficit set to 2 kPa. In order to test the influence of variation in leaf thickness on *g*_s, op_ we used three values of *d*_y_ (70, 100 and 130*μ*m).

### Analyses of trait evolution

To determine the temporal patterns of trait evolution, we generated a phylogeny from the list of taxa (Table S1) using Phylomatic (v. 3) and its stored family-level supertree (v. R20120829). To date nodes in the supertree, we compiled node ages from recent, fossil-calibrated estimates of crown group ages. Node ages were taken from Magallón et al. ^26^ for angiosperms, Lu et al. ^27^ for gymnosperms, and Testo and Sundue ^28^ for ferns. The age of all seed plants was taken as 330 million years ^29^. Because there is some uncertainty in the maximum age of the ancestor of all angiosperms, we took the angiosperm crown age used by Brodribb and Field ^30^ to make our results directly comparable to theirs. We tested this assumed angiosperm age by using different ages for the crown group angiosperms ranging from 130 Ma to 180 Ma, and the results were not qualitatively different. Of the 331 internal nodes in our tree, 90 of them had ages. These ages were assigned to nodes and all other branch lengths smoothed using the function ‘bladj’ in the software Phylocom (v. 4.2 ^31^). Polytomies were resolved by random bifurcation and adding 5 million years to each of these new branches and subtracting an equivalent amount from the descending branches so that the tree remained ultrametric. For all subsequent analyses of character evolution, this method for randomly resolving polytomies was repeated 100 times to account for phylogenetic uncertainty. To fit models of trait evolution, stochastic character change ^32^ was mapped on each randomly resolved tree using the function ‘make.simmap’ in the R package *phytools* ^33^ before fitting each model of evolution (described below). For ancestral state reconstructions the ages and character estimates at each node were averaged across the 100 randomly resolved trees.

Ancestral state reconstructions were calculated using the residual maximum likelihood method, implemented in the function ‘ace’ from the R package *ape* ^34^. To determine when changes in traits pushed the frontiers of trait values, the upper (*D*_v_) and lower (genome size and *l*_g_) limits of traits were estimated by first extracting the upper or lower ten percent of reconstructed trait values in sequential five million year windows and then attempting to fit curves to these values. This method is similar to a previous analysis of *D*_v_ evolution through time ^35^, which is included here for comparison. We compared three types of curve fits: a linear fit that lacked slope (equivalent to the mean of the reconstructed trait values), a linear fit that included both a slope and an intercept, and a nonlinear curve of the form *trait* = *a* + *b*/(1 + *e*^(-(*time* + *c*)/*d*)). Curves were fit to reconstructed trait values for each clade between 160 and 50 Ma, which corresponds to the time period encompassing the major diversification and expansion of the angiosperms, and the best fit was chosen based on AIC scores with a difference in AIC of 5 taken to indicate significant differences in fits. Ancestral state reconstructions of genome size for CAM angiosperms were calculated separately from C3 and C4 angiosperms because of the computational time required for the analyses. Phylogenetic independent contrasts (PICs) were used to determine whether traits underwent correlated evolution. PICs for each pairwise combination of traits were calculated for only species with data for both traits. Correlations between PICs were calculated using Spearman rank correlations in the function ‘cor.table’ from the R package *picante* ^36^.

To determine whether the tempo and mode of genome size evolution differed among major clades and lineages with different photosynthetic pathways, we used the R package *mvMORPH* ^37^ to fit four types of evolutionary models under a maximum likelihood criterion: Brownian motion (BM) with a single rate of evolution for the entire tree, Brownian motion with multiple rates for different groups of taxa, Ornstein-Uhlenbeck (OU) process with a single adaptive optimum for all species, Ornstein-Uhlenbeck process with different trait optima for different groups of taxa. Three types of regimes were modeled: (1) C3 species in all three major clades, (2) angiosperms differing in photosynthetic pathway, (3) all clades and all photosynthetic pathways. In all of these analyses, we accounted for phylogenetic uncertainty as described above. Model fits were compared using AIC scores with a difference in AIC of 5 assumed to indicate a significantly better model. In determining whether genome size evolution differed among angiosperms with different photosynthetic pathways, we attempted to account for the large discrepancy in the number of C3 and CAM angiosperms in the dataset by using all species (‘unbalanced’ analysis) and by randomly sampling 271 CAM species so that there were equivalent numbers of C3 and CAM species (‘balanced’ analysis). Then the same models as above were fit and compared. Because the analysis focused on the comparison between C3 and CAM species, *t*-tests were used to compare phylogenetic means and rates of genome size evolution, although estimated parameters for C4 species are included for completeness.

To determine whether there were temporal lags between changes in genome size and changes in *D*_v_ and *l*_g_, we compared OU models that allowed for multiple trait optima that used symmetric (no time lag) and non-symmetric (one trait lags behind another) alpha matrices. Although in univariate analyses the OU model underperformed the BM models, analyses of time lags between trait shifts can be assessed using only OU models.

### Scaling relationships

Scaling relationships between genome size and *D*_v_, *l*_g_, *g*_s,max_, and *g*_s,op_ were calculated from log-transformed data and analyzed using the function ‘sma’ in the R package *smatr* ^36^. Analyses were performed for the entire dataset and also for individual clades. Slope tests were used to determine whether the scaling relationship between genome size and *g*_s,max_ was significantly different than the relationship between genome size and *g*_s,op_ and whether the scaling relationship between genome size and *g*_s,op_ and *g*_s,max_ differed among clades.

---

**Table S2.** Trait and phylogenetic independent contrast (PIC) correlations for all species and for only the angiosperms. Trait correlations are in the upper triangle and contrast correlations in the lower triangle. Spearman rank correlation coefficients are shown. Asterisks indicate significance level: *P < 0.05; **P < 0.01; ***P < 0.001

| All species |  |  |  |  |  |
| --- | --- | --- | --- | --- | --- |
| | Genome size | $D_v$ | $l_g$ | $g_{s, max}$ | $g_{s, op 70}$ |
| Genome size |  | -0.65*** | 0.74*** | -0.46*** | -0.65*** |
| $D_v$ | -0.16* | | -0.63*** | 0.67*** | 0.99*** |
| $l_g$ | 0.46*** | -0.25* | | -0.31*** | -0.63*** |
| $g_{s, max}$ | -0.15* | 0.35** | 0.01 | | 0.67*** |
| $g_{s, op 70}$ | -0.19** | 0.99*** | -0.25* | 0.37** | |
| Angiosperms |  |  |  |  |  |
| Genome size |  | -0.48*** | 0.65*** | -0.34*** | -0.49*** |
| $D_v$ | -0.16* | | -0.56*** | 0.59*** | 0.99*** |
| $l_g$ | 0.45*** | -0.22 | | -0.23*** | -0.56*** |
| $g_{s, max}$ | -0.17* | 0.32* | 0.04 | | 0.59*** |
| $g_{s, op 70}$ | -0.19* | 0.99*** | -0.22 | 0.35** | |

